# Testing relationships between multiple regional features and biogeographic processes of speciation, extinction, and dispersal

**DOI:** 10.1101/2023.06.19.545613

**Authors:** Sarah K. Swiston, Michael J. Landis

## Abstract

The spatial and environmental features of regions where clades are evolving are expected to impact biogeographic processes such as speciation, extinction, and dispersal. Any number of regional features (such as altitude, distance, area, etc.) may be directly or indirectly related to these processes. For example, it may be that distances or differences in altitude or both may limit dispersal rates. However, it is difficult to disentangle which features are most strongly related to rates of different processes. Here, we present an extensible Multi-feature Feature-Informed GeoSSE (MultiFIG) model that allows for the simultaneous investigation of any number of regional features. MultiFIG provides a conceptual framework for incorporating large numbers of features of different types, including categorical, quantitative, within-region, and between-region features, along with a mathematical framework for translating those features into biogeographic rates for statistical hypothesis testing. Using traditional Bayesian parameter estimation and reversible-jump Markov chain Monte Carlo, MultiFIG allows for the exploration of models with different numbers and combinations of feature-effect parameters, and generates estimates for the strengths of relationships between each regional feature and core process. We validate this model with a simulation study covering a range of scenarios with different numbers of regions, tree sizes, and feature values. We also demonstrate the application of MultiFIG with an empirical case study of the South American lizard genus Liolaemus, investigating sixteen regional features related to area, distance, and altitude. Our results show two important feature-process relationships: a negative distance/dispersal relationship, and a negative area/extinction relationship. Interestingly, although speciation rates were found to be higher in Andean versus non-Andean regions, the model did not assign significance to Andean- or altitude-related parameters. These results highlight the need to consider multiple regional features in biogeographic hypothesis testing.

## Introduction

Biogeographic processes like speciation, extinction, and dispersal are expected to be shaped by the spatial context in which these processes operate. Any number of regional features – such as those relating to region size, distance, altitude, climate, barrier-presence, or status as an island or continent – may directly or indirectly shape the biogeography of a particular clade. For example, an inverse relationship between island size and probability of extinction is an essential feature of MacArthur and Wilson’s theory of island biogeography (Macarthur and Wilson 1967), and region area has since been frequently implicated to explain patterns of diversification (Connor and McCoy 1979), particularly when historical areas are considered (Fine and Ree 2006). Similarly, an inverse relationship between distance and dispersal has been established, both in terms of physical distance (Macarthur and Wilson 1967) and traversability of connecting habitat (Adriaensen et al. 2003). It has also been suggested that climatic features influence diversification dynamics in a wide variety of clades, including plants (Scheiner and Rey-Benayas 1994), mammals (Buckley et al. 2010), amphibians and reptiles (Navas 2002), and many others.

Many different phylogenetic models of historical biogeography have been created for the purposes of explaining how species both evolve over time and move through space (Ronquist 1997; Ree et al. 2005; Goldberg et al. 2011; Matzke 2014; Valente et al. 2015; Albert et al. 2017; Skeels and Cardillo 2018; Quintero and Landis 2020; Landis et al. 2021, 2022). These models vary widely in function and application. However, even with the quantity and breadth of available models, testing hypotheses about the relationships between regional features and evolutionary/biogeographic rates remains a challenge. Due to its inference capabilities and incorporation of state-dependent diversification, the GeoSSE model (Goldberg et al. 2011) provides a good foundation for testing regional hypotheses of historical biogeography (Ree and Sanmartín 2009, 2018; Crisp et al. 2011).

The GeoSSE model is based on previous state-dependent speciation and extinction (SSE) models, primarily the Binary State Speciation and Extinction (BiSSE) model (Maddison et al. 2007). Previous SSE models such as BiSSE were used for character evolution rather than biogeography, but the two subjects have many similarities (species range could be considered an evolving character). The GeoSSE framework, which has generally only been applied to biogeographic systems with two regions, assigns one free parameter to every possible speciation, extinction, and dispersal rate and, because of this, we may be able to link the features of those regions to evolutionary and biogeographic rates. Then, one can jointly examine diversification/biogeographic processes and the individual effects of features on those processes. Unfortunately, the number of free parameters in a standard GeoSSE analysis explodes as the number of regions increases, complicating how we test relationships between regional features and biogeographic event rates. The Feature-Informed GeoSSE (FIG) model (Landis et al. 2022) addresses this issue by modeling biogeographic event rates as parametric functions of regional features. This parameterization strategy both collapses model complexity into something manageable while also creating a framework to test hypotheses about the importance of certain features on evolutionary processes. For example, the original FIG study found that distance and water barriers decreased dispersal rates and increased between-region speciation rates in *Anolis* lizards, as one might expect.

Because of the number of possible regional features (abiotic and biotic) that might influence biogeographic rates, it can be difficult to determine which features impact which processes. This is a major obstacle for hypothesis testing, since the quality of our phylogenetic inferences about the past (biogeographic rates and ancestral states) rely on our ability to choose the best-fitting models. Statistical model selection in historical biogeography has been useful for a wide variety of problems (Landis et al. 2013; Matzke 2014; Cardillo et al. 2017; Landis et al. 2022; Quintero et al. 2023), but typically, the procedure only compares models from a small space of low-complexity models. For example, it could compare a model that includes a distance-dependent dispersal rate parameter with a model that does not. However, we expect that a large number of regional features may influence dispersal rates, such as distances, water-barriers, altitudinal differences, climatic differences, etc. Testing for the effects of all combinations of features on evolutionary rates is challenging because of the vast model space that must be explored – roughly 2^*N*^ model comparisons for *N* possible feature-process relationship parameters that can be ‘on’ or ‘off’.

Bayesian phylogenetic methods have often overcome the problem of finding sets of good-fitting models using reversible-jump Markov chain Monte Carlo (RJMCMC) (Green 2003; Huelsenbeck et al. 2004). One of the simplest RJMCMC algorithms allows the Markov chain to move among a family of nested models by turning certain parameters ‘on’ or ‘off’, while also exploring parameter space like a traditional MCMC. Landis et al. (2022) used RJMCMC to fit FIG models to simulated data and an empirical dataset for *Anolis* to control model complexity and to test combinations of biogeographic hypotheses involving just four regional features: region size, distance, water-barrier presence, and insular region status.

In this paper, we introduce MultiFIG, a scalable Bayesian phylogenetic framework to efficiently explore a large set of biogeographic models for which any number of regional features may or may not change different biogeographic rates of speciation, extinction, and/or dispersal. MultiFIG builds upon the previous FIG model by allowing for the inclusion of any number of quantitative and categorical features without requiring their interactions to be predetermined. We then apply MultiFIG to a variety of historical biogeographic scenarios to characterise the circumstances for which our hypothesis testing framework performs well or poorly. These scenarios vary in the number of regions, number of taxa, values and variances of geographic features, and strength/direction of feature effects on evolutionary processes. By examining this breadth of simulation scenarios, we are able to assess model performance under a variety of conditions that may resemble different biological systems where taxa are evolving.

Finally, as a proof-of-concept, we apply the MultiFIG model to an empirical study system: *Liolaemus. Liolaemus* is a diverse genus of 199 lizard species that occur primarily within the central and southern Andean highland and the flanking lowland regions. Their biogeography has been investigated previously using a GeoSSE model with two regions (Andean versus non-Andean), although the existing model was not designed to untangle the effects of many regional features on diversification and dispersal (Esquerré et al. 2019). Therefore, we apply MultiFIG to *Liolaemus*, demonstrating how our framework can uncover more refined relationships between regional features and biogeographic rates, and how that refinement may influence historical biogeographic estimates.

## Methods

Here, we describe an extensible multi-feature Feature-Informed GeoSSE (MultiFIG) model. We perform a simulation experiment to validate the model and explore its behavior under different conditions, focusing on different values and variances of geographic features, numbers of regions, and numbers of taxa. Then, we apply the model to investigate how regional features related to area, distance, and altitude have impacted the evolution and historical biogeography of *Liolaemus*.

### Model definition

#### Discrete biogeographic ranges

The Feature-Informed GeoSSE (FIG) model (Landis et al. 2022) is an extension of the GeoSSE model (Goldberg et al. 2011), which is a member of a class of state-dependent speciation and extinction (SSE) models (Maddison et al. 2007). SSE models assign lineages to certain ‘states’ that correspond to some attribute they possess, such as a morphological character or a region of residence. Based on the state of a lineage, it experiences different rates of speciation and extinction, as well as rates of transition to other states. The GeoSSE model specifically handles cases where the states represent possible geographic ranges, comprised of one or more discrete regions inhabited by a species. Take, for example, a clade evolving among *N* = 3 discrete regions (*A, B*, and *C*). The subset of regions where a lineage is present defines its range, and its ‘state’ within the SSE framework. In the case of a 3-region system, a species can occupy any of 2^*N*^ *−* 1 = 7 possible geographic states at a given time: : *{A}, {B}, {C}, {A, B}, {A, C}, {B, C}*, or *{A, B, C}*. Species transition between these states through the processes of dispersal, local extinction (extirpation), and speciation (cladogenesis). Cladogenetic events may directly result in range changes (i.e. daughter lineages do not necessarily possess the same state as the parent). Therefore, the model incorporates four core processes of diversification through space and time: within-region speciation, between-region speciation, extirpation, and dispersal (Figure 1 depicts GeoSSE dynamics in a 2-region system).

**Figure 1:**
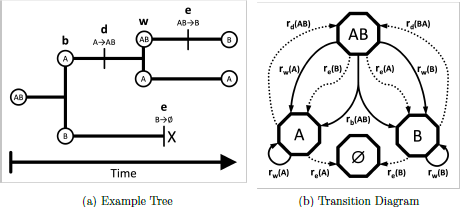
(a) Example phylogeny showing all four GeoSSE event types involving a biogeo-graphic system with two regions (A and B): within-region speciation [w], between-region speciation [b], extinction [e], and dispersal [d]. (b) Transition diagram showing event types associated with transitions between different states in a two-region system (GeoSSE model), following the structure of Goldberg et al. (2011) but using FIG notation. Cladogenetic events are shown using solid arrows, while anagenetic events are shown with dashed arrows.

#### Model events and rates

The model is constrained such that a lineage can only experience a single event at any instant in time. Dispersal and extirpation are anagenetic processes, operating between cladogenesis events along the branches of an evolutionary tree. For example, dispersal from *A* to *C* may cause an original range to expand from *{A, B}* to *{A, B, C}*. This allows us to define rates of dispersal events *r*_*d*_(*i, j*) for each pair of regions, which can be summed to produce rates of dispersal from any currently inhabited region into a new region. The extirpation rate *r*_*e*_(*i*) represents the rate at which a species loses region *i* from its range. A lineage experiences global extinction when it is extirpated from the final occupied region in its range. For example, extirpation in region *A* may cause an original range to shrink from *{A, B}* to *{B}*. If that species was then extirpated in region *B*, the lineage would be globally extinct and no longer diversify or disperse.

Within- and between-region speciation are cladogenetic processes, describing different range-inheritance scenarios at the nodes of an evolutionary tree. Let *R* be the ancestral range, and let *S* and *T* be the ranges of the two daughter lineages. Within-region speciation occurs within a single region in a species range, producing one daughter that inherits the parent’s entire range and one daughter with a range consisting of a single region. In a within-region speciation event, *R* = *S* and *T ∈ S*. For example, an ancestral range *R* = *{A, B}* may yield species with ranges *S* = *{A, B}* and *T* = *{A}*. Note that both *S* and *T* will always be equal to *R* when the ancestral range consists of only one region. We define rates of within-region speciation events *r*_*w*_(*i*) for each region *i*. For lineages with widespread ranges comprised of multiple regions, a speciation event may also occur between regions, producing two daughters that inherit non-overlapping subsets of the parent’s range. This event is only valid for ancestral ranges consisting of two or more regions. In a between-region speciation event, *R* = *S ∪ T* and *S ∩ T* = *∅*. For example, an ancestral range *R* = *{A, B, C}* may split into daughters *S* = *{A, B}* and *T* = *{C}*. This allows us to define rates of between-region speciation events *r*_*b*_(*S, T*), which are related to sets of region pairs (*i, j*) through a range split score to prevent explosive model overparameterization (Landis et al. 2022).

The parameters associated with these four processes are functions of individual regions, so the evolutionary rates experienced by a lineage are functions of the regions where that lineage is present (and their features). This results in four rate parameters for each region *i* or region pair *i, j*: *r*_*e*_(*i*) for extinction, *r*_*w*_(*i*) for within-region speciation, *r*_*d*_(*i, j*) for dispersal, and *r*_*b*_(*S, T*) for between-region speciation.

#### MultiFIG rates

In the basic GeoSSE model, all regions or region pairs are assigned their own independent parameters representing the core processes, which are dissociated from any regional features that may be causing or correlated with rates variation. This is tractable for a scenario with few regions (the creators of the GeoSSE model applied a system with *N* = 2 regions), but it presents several problems when the number of regions is large. Exploring a large state space (all possible region combinations) makes it challenging to calculate likelihoods, causing performance degradation. We use TensorPhylo to address this issue (May and Meyer 2022). Also, estimating so many distinct rates means estimating large numbers of distinct event rate parameters (see Landis et al. (2022) Supplementary Table S2). Because of this, we may not have sufficient data to estimate these parameters with any accuracy. Another problem is that large numbers of free parameters create a highly multimodal posterior surface, which means that Bayesian inferences like MCMC may mix slowly, leading to high computational costs. This leaves many questions with large state spaces and/or complex biogeographic model structures understudied. However, if we expect that certain features shape biogeographic rates in similar ways across regions, we can express evolutionary rates as functions of the features of the regions where lineages are evolving. For instance, we may hypothesize that dispersal between two regions scales with distance, and we can therefore represent dispersal as a function of the data (known distances between regions) and a single parameter describing the strength and direction of the effect of distance on dispersal. This allows us to reduce the number of model parameters and to directly examine whether particular region features correlate with increased or decreased rates of certain processes. The Feature-Informed GeoSSE (or FIG) model (Landis et al. 2022) presents a way to parameterize the combined effects of many region features, allowing for inference of biogeographic history, evolutionary rates, and model parameters in scenarios with numerous regions.

The FIG model incorporates geographical features with two value types as model variables: quantitative features and categorical features. Quantitative features have continuous real values while categorical features have discrete values. In our analyses, we only used positive-valued quantitative features and binary-valued categorical features, though these assumptions can be relaxed to suit the needs of the research question. The two value types are handled differently by the model, but are ultimately transformed so that they can collectively contribute to evolutionary rates. The MultiFIG model also separates data by dimensionality type. Data can represent one-dimensional within-region information (values for each individual region) or two-dimensional between-region information (values for each region pair). We use four different containers to represent these data types. Each of these containers can store data for several different features layers of a given type that are indexed by *k* (Figure 2).

**Figure 2:**
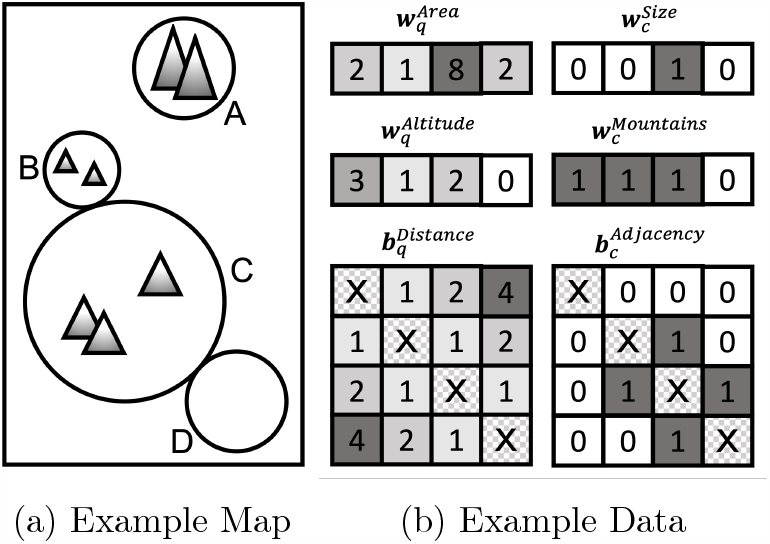
(a) A visual representation of a hypothetical biogeographic system with four regions (A, B, C, D) with different quantitative (area, altitude, distance) and categorical features (size, mountain-presence, adjacency). (b) Regional features of the biogeographic system are encoded by MultiFIG into variables containing quantitative data (numerical values in **w**_*q*_ and **b**_*q*_), categorical data (binary values in **w**_*c*_ and **b**_*c*_), within-region data (vectors **w**_*c*_ and **w**_*q*_), and between-region data (matrices **b**_*c*_ and **b**_*q*_).

Quantitative feature data are stored in the *N* -vector 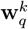 for within-region features and in the *N × N* -matrix 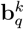 for between-region features, where *N* is the number of regions in the system. For instance, 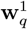 might contain areas (e.g. km^2^) for each region, while 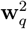 may contain mean altitudes for each region. Vectors in **w**_*c*_ and matrices in **b**_*c*_ are used similarly to represent data about within- and between-region categorical data. Note that the feature layer indexed by a specific value of *k* in one container does not necessarily imply anything about the layer indexed by the same value of *k* in another container – e.g. layers indexed by *k* = 2 do not all necessarily describe features associated with altitude. For instance, 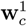 might contain a vector categorizing regions as ‘small/large’, and 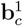 might contain a matrix representing region pair adjacency/non-adjacency. The processes of extinction and within-region speciation make use of within-region data, while the processes of dispersal and between-region speciation make use of between-region data. Table 1 describes the different features used in our simulation study and empirical analysis, giving an example of the types of data that may be included in these different containers. However, the framework is general, and features can be added or substituted as needed for different study systems.

**Table 1:** Overview of relationships between each feature and the associated MultiFIG parameters. Each feature (**W** or **b**) is associated with two of four processes: within-region speciation (w), extinction (e), between-region speciation (b), and dispersal (d). The strength of the effect is controlled through a set of estimated parameters (*ϕ* or *σ*). Details provided in the main text.

| Feature | Description | Processes | Parameters |
| --- | --- | --- | --- |
| $\mathbf{w}_q^{Area}$ | areas of each region | w, e | $\phi_w^{Area}, \phi_e^{Area}$ |
| $\mathbf{w}_q^{Altitude}$ | altitudes of each region | w, e | $\phi_w^{Altitude}, \phi_e^{Altitude}$ |
| $\mathbf{b}_q^{Distance}$ | mean distances between each region pair | b, d | $\phi_b^{Distance}, \phi_d^{Distance}$ |
| $\mathbf{b}_q^{Altitude}$ | differences in mean altitude between each region pair | b, d | $\phi_b^{Altitude}, \phi_d^{Altitude}$ |
| $\mathbf{w}_c^{Area}$ | sizes of each region (large/small) | w, e | $\sigma_w^{Area}, \sigma_e^{Area}$ |
| $\mathbf{w}_c^{Altitude}$ | heights of each region (highland/lowland) | w, e | $\sigma_w^{Altitude}, \sigma_e^{Altitude}$ |
| $\mathbf{b}_c^{Distance}$ | adjacency of each region pair (adjacent/non-adjacent) | b, d | $\sigma_b^{Distance}, \sigma_d^{Distance}$ |
| $\mathbf{b}_c^{Altitude}$ | comparison of heights of each region pair (same/different) | b, d | $\sigma_b^{Altitude}, \sigma_d^{Altitude}$ |

The MultiFIG model associates regional feature data with biogeographic rates through a series of parameterized transformations (Figure 3). The function 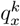 is used to handle quantitative data, and parameterizes the effects of a quantitative feature layer *k* on the biogeographic process *x*, where *x* can represent dispersal (*d*), extinction (*e*), within-region speciation (*w*), or between-region speciation (*b*). Each quantitative feature layer *k* is assigned a single set of 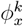 parameters. These parameters represent the strength and direction of the effect of each feature on a particular process (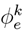 for extinction, 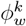 for within-region speciation, 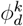 for dispersal, and 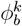 for between-region speciation). All regions or region pairs share one 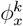 per feature and process. For each feature layer *k* and region *i* (or region pair *i, j*), the value of the feature for the region/pair is normalized by the geometric mean of the feature over all regions/pairs, *gm*(·), then exponentiated by the effect parameter 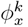. Thus, the vector 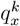 represents the effects of each quantitative feature *k* on a particular process *x*, and contains information about each region *i* or region pair *i, j*.

**Figure 3:**
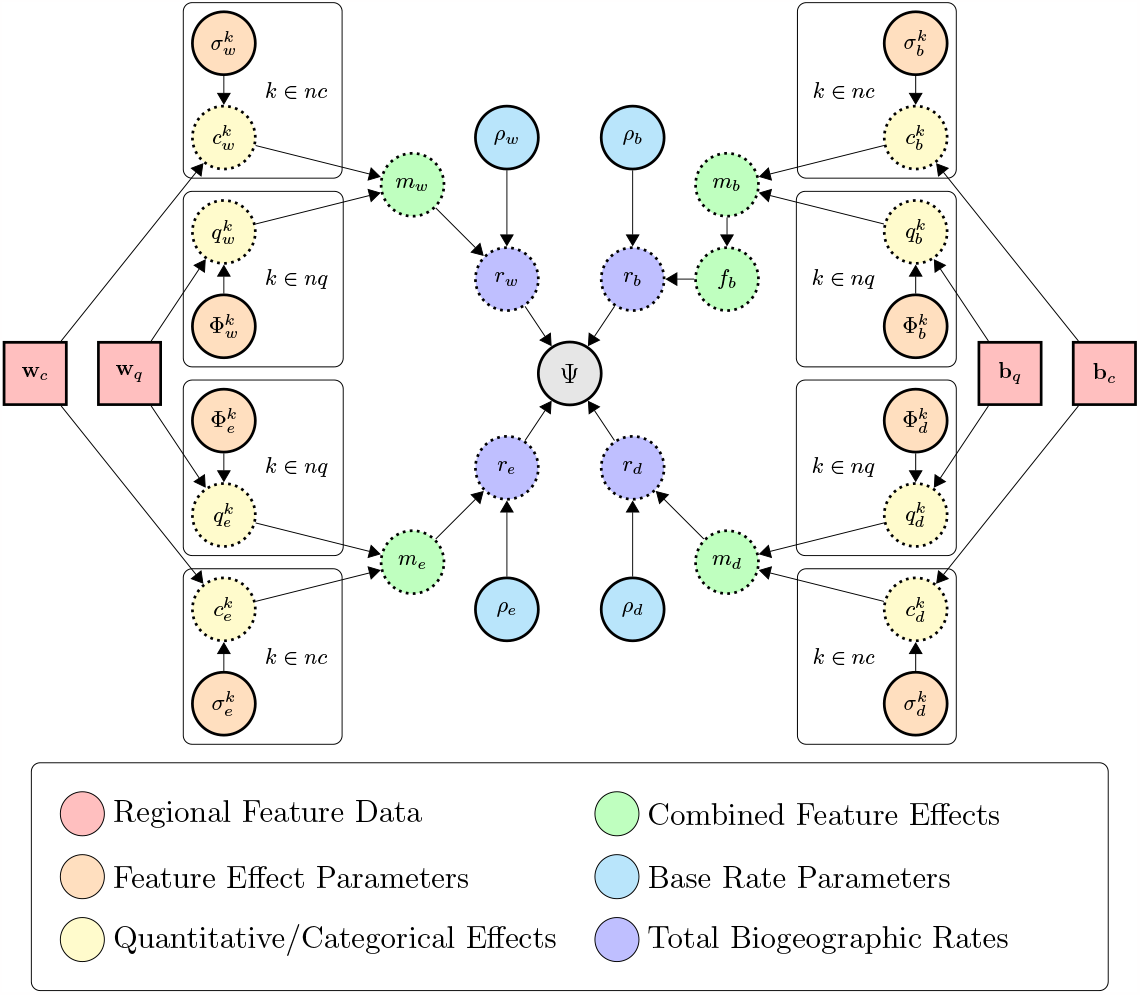
Full graphical model for generalized MultiFIG. The main text and Table describe how to interpret regional feature data (**w, b**; red), feature effect parameters (**Φ**, *σ*; orange), individual feature effects (*q, c*; yellow), combined feature effects (*m*) and the split score function (*f*_*b*_; green), base rates (*ρ*; cyan), the absolute biogeographic rates (*r*; blue), and the GeoSSE process (Ψ; gray). Graphical model notation follows the conventions described in Höhna et al. (2014). Square nodes represent constant values (data). Circle nodes with solid lines represent stochastic variables (model parameters, and the phylogeny, which is fixed in this analysis). Circle nodes with dotted lines represent deterministic variables (functions). Large rectangles indicate iterative plates.

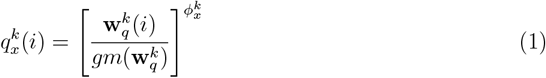

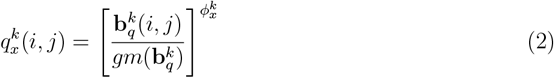

Similarly, the function 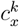 is used to handle categorical data, and parameterizes the effects of a categorical feature layer *k* on process *x*. Each categorical feature layer *k* is assigned a single set of 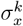 parameters. These parameters represent the strength and direction of the effect of each feature on a particular process (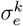 for extinction, 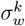 for within-region speciation, 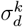 for dispersal, and 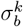 for between-region speciation). All regions or region pairs share one 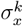 per feature and process. For each feature layer *k* and region *i* (or region pair *i, j*), an indicator function is used to determine which category the region/pair falls into (here labeled *A* and *B*). If the region/pair is in category *A*, the indicator function produces 0, and if it is in state *B*, the indicator function produces 1. This indicator function is used to exponentiate the 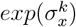 parameter, which means the 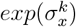 function will output 1 if the region/pair is in category *A*, and 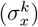 if the region/pair is in category *B*. Thus, the vector 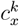 represents the effects of each categorical feature *k* on a particular process *x*, and contains information about each region *i* or region pair *i, j*.

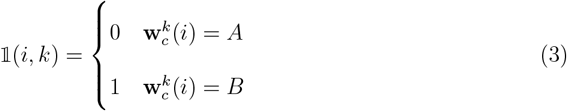

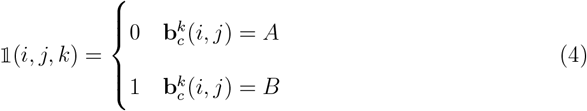

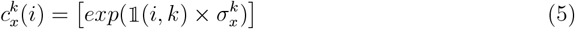

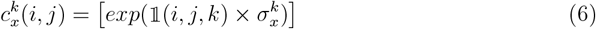

The form of this equation (making use of an indicator function) is more general than necessary for the analyses presented here, since we represented all categorical data according to a 0/1 binary. However, it displays how the framework could be adapted for more complicated sets of possible categorical values.

The *m*_*x*_ function combines the effects of all categorical and quantitative feature layers for a given region *i* or region pair *i, j*. Each process has its own *m*_*x*_ vector containing information about the total effects of all features on each region or region pair (*m*_*e*_(*i*), *m*_*w*_(*i*), *m*_*d*_(*i, j*), *m*_*b*_(*i, j*)). Note that parameter values of 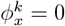 or 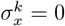 yield 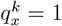 and 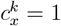 for all regions *i* or region pairs *i, j*, indicating that the feature will not influence region-specific event rates (a null effect).

Relative rate of extinction or within-region speciation in region *i* are computed as the product over all relative rate contributions for all relevant quantitative and categorical effects for region *i*, with the general form

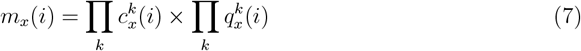

whereas analogous dispersal and between-region speciation rate modifiers involving regions *i* and *j* have the form

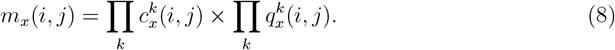

Finally, the *m*_*x*_ values are scaled by a base rate parameter for each process, *ρ*_*x*_. As before, the absolute extinction or within-region speciation rate for region *i* is computed as

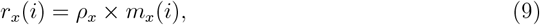

the absolute dispersal rate between regions *i* and *j* is computed as

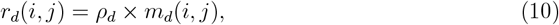

while the absolute between-region speciation rate for splitting range *R* into daughter ranges *S* and *T* is

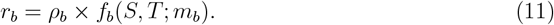

where *f*_*b*_ is the range split score function, which is used to calculate the relative rates of between-region speciation for different splits (Landis et al. 2022), and is defined as

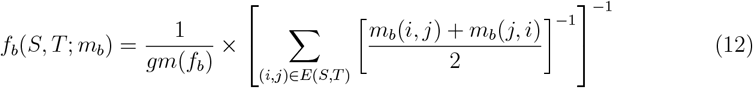

In effect, *f*_*b*_ assigns smaller values to ranges that are more ‘cohesive’, where cohesiveness is measured by how easily the current range *R* = *S ∪ T* can be split into subranges *S* and *T* given the strength of connectivity between regions modeled by *m*_*b*_. Let the original range be a fully connected graph, while treating occupied regions in *R* = *S ∪ T* as nodes and the matrix *m*_*b*_ as edge weights. *E*(*S, T*) represents the cutset, which is the minimal set of edges that must be cut to fully separate the nodes in range *S* from the nodes in range *T*.

This produces a set of absolute process rates *r*_*e*_(*i*), *r*_*w*_(*i*), *r*_*d*_(*i, j*), and *r*_*b*_(*S, T*) for all regions in the biogeographic system, representing the rates of extinction, within-region speciation, dispersal, and between-region speciation in the GeoSSE model. However, unlike in the GeoSSE framework, every distinct biogeographic event is not assigned its own free rate parameter; they are instead modeled using observable data and a much smaller set of parameters that remains constant with regard to the number of regions in the system (Figure 3).

### Model selection

The MultiFIG modeling framework allows for the inclusion of numerous regional features that may or may not correlate with evolutionary rates. Typically, determining which parameters are or are not necessary in a model requires one to test each possible combination of parameters, and choose the best model according to some criteria. For example, one could use Bayes factors computed using marginal likelihoods to compare models, requiring independent MCMC runs for each model being compared. However, using reversible-jump Markov chain Monte Carlo (RJMCMC), it is possible to ‘jump’ between models with different dimensionality (Green 2003). Because of this, RJMCMC allows us to perform model selection in addition to parameter estimation. In practice, a new proposal is added to the MCMC for each feature effect parameter *ϕ* or *σ*, which suggests that the parameter’s value is actually equal to 0, effectively turning the parameter ‘off’. The reverse proposal is also added, turning the parameter back ‘on’ and allowing the MCMC to continue to explore non-zero parameter values. For simple nested models, the Metropolis-Hastings-Green ratio for transdimensional proposals is very straightforward to compute, assuming that the priors distributing reversible-jump variables assume independence (Green 1995). RevBayes allows for the implementation of reversible-jump MCMC moves (Höhna et al. 2019). We can keep track of the amount of time the Markov chain spends with a parameter ‘on’ versus ‘off’, which can be used to obtain a posterior probability. In this way, we are able to examine whether various features have an effect on a particular process, or whether they have no effect (an effect parameter value of 0). Note that the term ‘effect’ here refers to a mathematical relationship, and does not necessarily indicate causality; a feature with an inferred effect may not have contributed to diversification or biogeographic rates directly. Rather, it may be correlated with some other, unexamined feature that influenced these processes.

Since geographical features themselves are often correlated, especially when both quantitative and categorical versions of the same feature are incorporated, it is essential that we are able to interpret reversible-jump posteriors with respect to the ‘on/off’ status of other parameters. Fortunately, RJMCMC results allow us to examine the joint probabilities of different parameter combinations. For example, we might obtain the following diagram representing joint probabilities for the effect/non-effect of two parameters connected to closely-related phenomena, such as highland/lowland (*σ*) and mean altitude (*ϕ*).

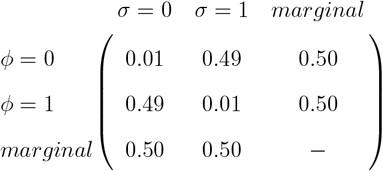

Marginal probabilities represent the probability of an event independent of other events, and are typically returned by MCMC analyses. By examining the marginal probabilities associated with each parameter in the above example, it could not be concluded that either parameter has an effect. For instance, the marginal probability of *ϕ* = 1 is 0.50, and the marginal probability of *ϕ* = 0 is 0.50. However, by examining the joint probabilities, we find that the scenarios where one of the parameters has an effect are much more probable than the scenario where neither parameter has an effect. The probability of *ϕ* = 1, *σ* = 0 is 0.49, the probability of *ϕ* = 0, *σ* = 1 is 0.49, and the probability of *ϕ* = 1, *σ* = 1 is 0.01, for a total probability of 0.99. The probability of *ϕ* = 0, *σ* = 0 is 0.01. Therefore, we could conclude that altitude, either in a categorical or quantitative context, likely has an effect on a certain process, even if the specifics (categorical or quantitative) are confounded by the model design.

### Simulation experiment

To assess the performance of our model, we simulated 900 biogeographic datasets under a range of scenarios that varied in terms of the number of simulated taxa, the number of regions, the values of regional features, and the values of evolutionary process parameters.

Using R, we simulated 450 different geographic scenarios with 3, 5, or 7 regions (150 geographies each). Each region/region pair within each geographic scenario was assigned a value for eight different region features. Four were quantitative: area, mean altitude, difference in mean altitude, and distance. The other four were categorical representations of the same quantitative features. Since all region features would later be normalized, the scales of the features were not selected to match a particular biological scenario. However, the variances of the features were designed such that the *Liolaemus* study system of Esquerré et al. (2019) would fall comfortably within the range of simulation scenarios tested (Figure 4).

**Figure 4:**
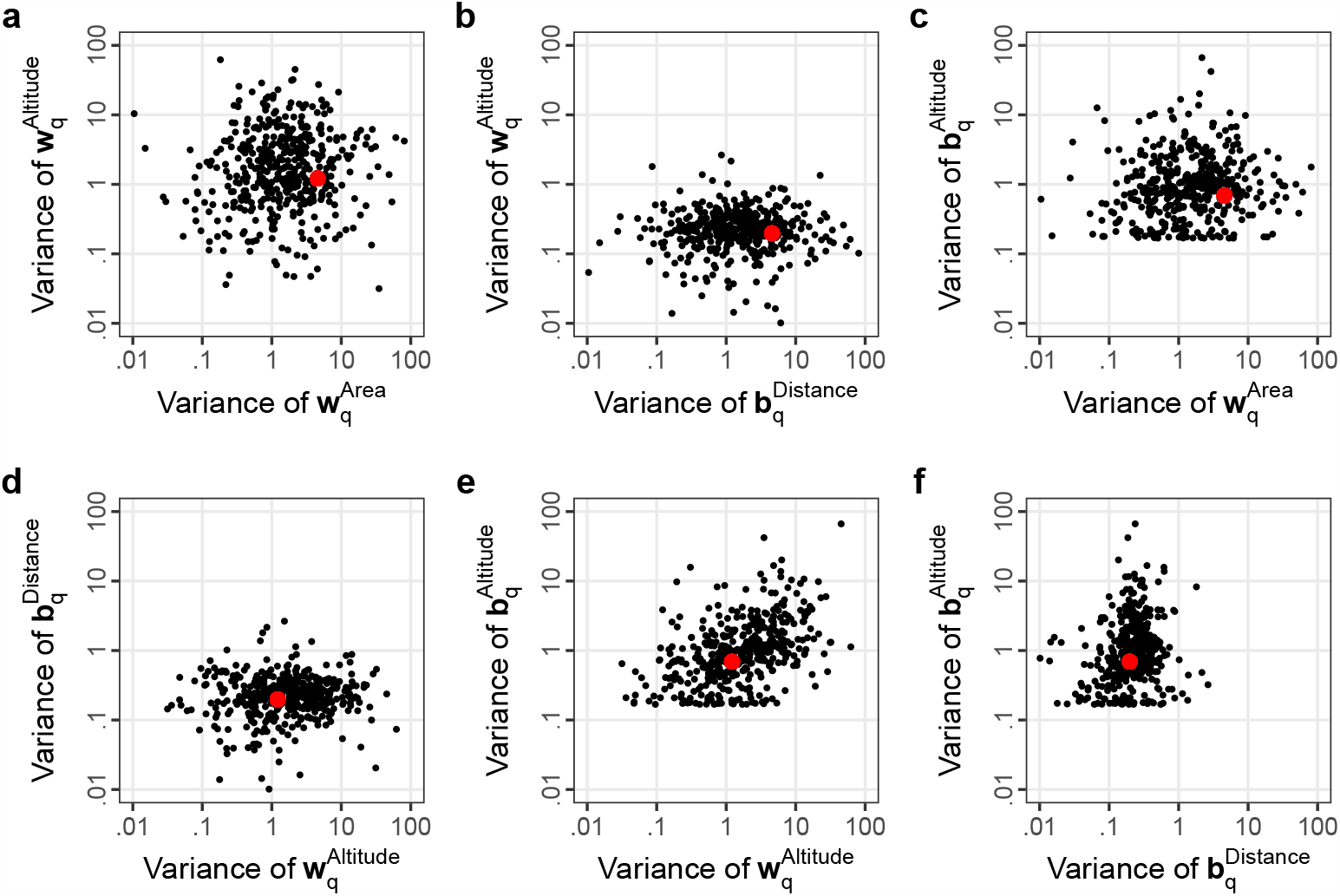
Pairwise variances of geographic features for 450 simulated geographies. Variances of regional features among the six South American regions used in our *Liolaemus* MultiFIG analysis are also included (red dots). Simulated scenarios cover a wide range of feature variances, and the variances of the South American regional features are in well-explored regions of parameter space.

Region areas were drawn randomly from a gamma distribution with shape 1 and rate 0.15. Categorical areas were then assigned to regions based on whether a region’s area was greater than (1) or less than (0) the mean area of all regions in that geographic scenario. Mean altitudes were drawn from the same gamma distribution, and categorical altitudes were assigned in the same manner. Quantitative differences altitude were calculated by taking pairwise differences between each region. Categorical differences in altitude were assigned based on whether each pair of regions had the same categorical state (1) or different states (0). To preserve distance properties, each region was assigned a ‘location’ consisting of *x* and *y* coordinates drawn from a uniform distribution on [0, 1]. Euclidean distances were then calculated between each region pair. Categorical distances (adjacency) were not calculated based on quantitative distances, as this does not always represent physical reality (as in the *Liolaemus* study system). Rather, region pairs were randomly assigned adjacency (1) or non-adjacency (0) with equal probability.

From the simulated geographies, we used RevBayes (Höhna et al. 2016; W. A. Freyman and S. Höhna 2017) to generate 900 simulated time-calibrated phylogenies and biogeographies. Using the 150 geographies with 5 regions, we created 600 phylogenies/biogeographies, with 150 simulations targeted at each tree size category: extra small (25–50 taxa), small (51–100 taxa), medium (101–200 taxa), and large (201–350 taxa). The number of taxa targeted by each simulation was drawn from a uniform distribution over the range for the size category. The other 300 geographies (150 with 3 regions and 150 with 7 regions) were used to simulate 300 phylogenies/biogeographies with medium-sized trees.

Simulations were performed under the MultiFIG model using RevBayes. All within-region features were modeled to act upon both within-region processes (extirpation and within-region speciation), whereas all between-region features were modeled to act upon both between-region processes (dispersal and between-region speciation). Feature effect parameters were drawn from a normal distribution with mean 0 and variance 1, while base rate parameters for each process were drawn from an exponential distribution with a rate of 1. A reversible-jump model was not used during simulations (no feature effect parameters were given a zero value), only for one set of inferences.

The simulations resulted in 900 unique phylogenies and sets of tip states (ranges of extant taxa). Tree heights varied, with a mean height of 2.8 and a standard deviation of 11.1. Of the simulated trees, 846 were adequate for inference (namely, no events happened nearly-concurrently within an extremely short [1e-9] interval of time). Using RevBayes (Höhna et al. 2016) and Tensorphylo (May and Meyer 2022), these simulated phylogenies/biogeographies were then analyzed under the generative model using both MCMC and RJMCMC (with equal probabilities assigned to zero and non-zero parameter values), to determine model performance in different regions of parameter space. MCMC and RJMCMC analyses were conducted over 10,000 generations, discarding 10% as burn-in (without tuning hyperparameters). Initially, we performed analysis of the simulated datasets using RJMCMC alone. The utility of Reversible Jump is primarily to select between models with different numbers of parameters, and given that we did not simulate under models with ‘off’ parameters, the coverage analysis indicated a mismatch between simulation and inference priors. This was expected. Therefore, to validate the model and assess the accuracy of parameter estimates, we did not use Reversible Jump for the coverage and percent error analyses, choosing to analyze the datasets with traditional MCMC for these purposes. We also performed standard MCMC on the *Liolaemus* study system to generate parameter estimates, in conjunction with the RJMCMC model selection results.

Using R, we summarized the Bayesian coverage of each model parameter, including base rates *ρ* for each process and feature-associated *σ* and *ϕ* parameters. For the simulation study only, we utilized 80% highest posterior density (HPD) intervals to calculate coverages. We chose to use HPD80 because HPD95 is expensive to estimate accurately, and the goal of the coverage analysis is to determine model performance, not to infer significance of particular parameters. If the inference machinery is behaving properly, and the model is correctly specified, we would expect a coverage value of 80% (80% of 80% HPD intervals should cover the truth). We also calculated median absolute percent error for different scenarios, indicating the expected accuracy of analyses fitting certain conditions.

### Liolaemus analysis

We applied our FIG model to an empirical study system, the *Liolaemus* lizards of South America, in order to demonstrate how our framework can reveal relationships between region features and evolutionary rates. We split the southern part of the continent of South America into 6 discrete regions for the MultiFIG analysis, as outlined in Esquerré et al. (2019), and generated spatial polygon representations of these regions using QGIS. Altitudinal data was obtained from the NASA Shuttle Radar Topographic Mission (NASA 2013) using the ‘raster’ package in R. We extracted data for 8 regional features (Figure 1): quantitative area 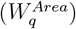, categorical area (above/below mean, 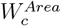), quantitative distance (mean distance between a randomly selected point in the first region and a randomly selected point in the second region, 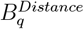), categorical distance (adjacent/non-adjacent, 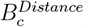), quantitative altitude (mean altitude, 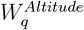), categorical altitude (above/below mean, 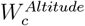), quantitative altitudinal difference (difference in mean altitude, 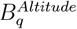), and categorical altitudinal difference (whether regions share highland/lowland status or not, 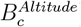).

The time-calibrated phylogeny of *Liolaemus* and present-day ranges were obtained from Esquerré et al. (2019). While the Esquerré et al. dataset includes data for the entire Liolaemidae family, we focus on the genus *Liolaemus*. The other two genera are older, less species-rich, and more narrowly distributed than *Liolaemus. Ctenoblepharys* is monotypic, and *Phymaturus* predominantly consists of Patagonian and Central Andean species. Therefore, to avoid the potential biases that these genera could introduced due to model misspecification, we exclude *Ctenoblepharys* and *Phymaturus*. We fixed the phylogeny of in this analysis, as there seemed to be no important effect of uncertainty on rate estimates under a time-constant FIG model with *Anolis* lizards (Landis et al. 2022). To infer rates of evolutionary processes and effect parameters associated with each feature, we used RevBayes (Höhna et al. 2016) to perform both standard MCMC and RJMCMC. For the standard MCMC, we ran the analysis for 10,000 generations with an additional 10% burn-in period to tune hyperparameters. For the RJMCMC, which struggled more with mixing, we ran the analysis for 20,000 generations with an additional 10% burn-in period to tune hyperparameters. Several independent runs were compared to assess consistency, but the results presented here are drawn from one standard MCMC analysis (posterior distributions) and one RJMCMC analysis (RJMCMC probabilities), as all analyses showed the same patterns. Priors were identical to those in the simulation experiment.

We analyzed the results using R, and determined which *σ* and *ϕ* parameters indicated strong effects (significant probability of nonzero value in RJMCMC, zero value outside 95% highest posterior density interval in standard MCMC). We also estimated base rates of each process (Figure S10) and generated an ancestral state tree using the MCMC output (Figure S15).

### MultiFIG code and tutorial

Raw data, R scripts, and RevBayes scripts used in these analyses are available in the code repository: https://bitbucket.org/sswiston/multifig/src/master/. The pre-configured Docker container used to run RevBayes MultiFIG scripts is available on Docker Hub: https://hub.docker.com/r/sswiston/rb_tp. The RevBayes website hosts a tutorial describing how to design a MultiFIG analysis for other clades and biogeographic systems: https://revbayes.github.io/tutorials/multifig.html.

## Results

### Simulation experiment

Of the 846 simulated datasets, 748 produced successful MCMC runs and 720 produced successful RJMCMC runs (87% success overall). Only 1 run encountered an error. The remaining MCMC and RJMCMC runs exceeded the maximum cluster wall time or encountered apparent computational difficulties during the analysis (e.g. MCMC getting ‘stuck’ in parameter space). Because these runs did not reach 10,000 generations, they were excluded from the analysis. Coverages were calculated for each model parameter, representing the number of datasets where the 80% HPD interval of the estimate contains the ‘true’ value of the parameter (the value used in simulation). A coverage analysis of the 634 simulated datasets indicates that the inference machinery behaves as expected (Table S2). However, mean coverage (77%) is slightly below the 80% benchmark. This could possibly be due to the way in which trees were generated. By targeting specific numbers of taxa (to ensure an even spread of tree sizes), many trees were rejected. Also, on occasion, the parameter values drawn from the prior were unable to create a valid tree after many attempts, and new parameter values had to be selected. Therefore, the true distribution of simulating parameters may not precisely match the priors used for simulation, causing a slight decrease in coverage due to misspecification. This does not indicate a problem with the model itself.

MultiFIG estimates evolutionary rates and feature effect parameters with a reasonable degree of accuracy (Figure 5). Median absolute percent errors give a more detailed view of the model’s overall performance (Figure S8). Generally, the model produced more accurate estimates of base rates in scenarios with larger trees, and it produced more accurate estimates of effect parameters (*ϕ* and *σ*) in scenarios with more regions.

**Figure 5:**
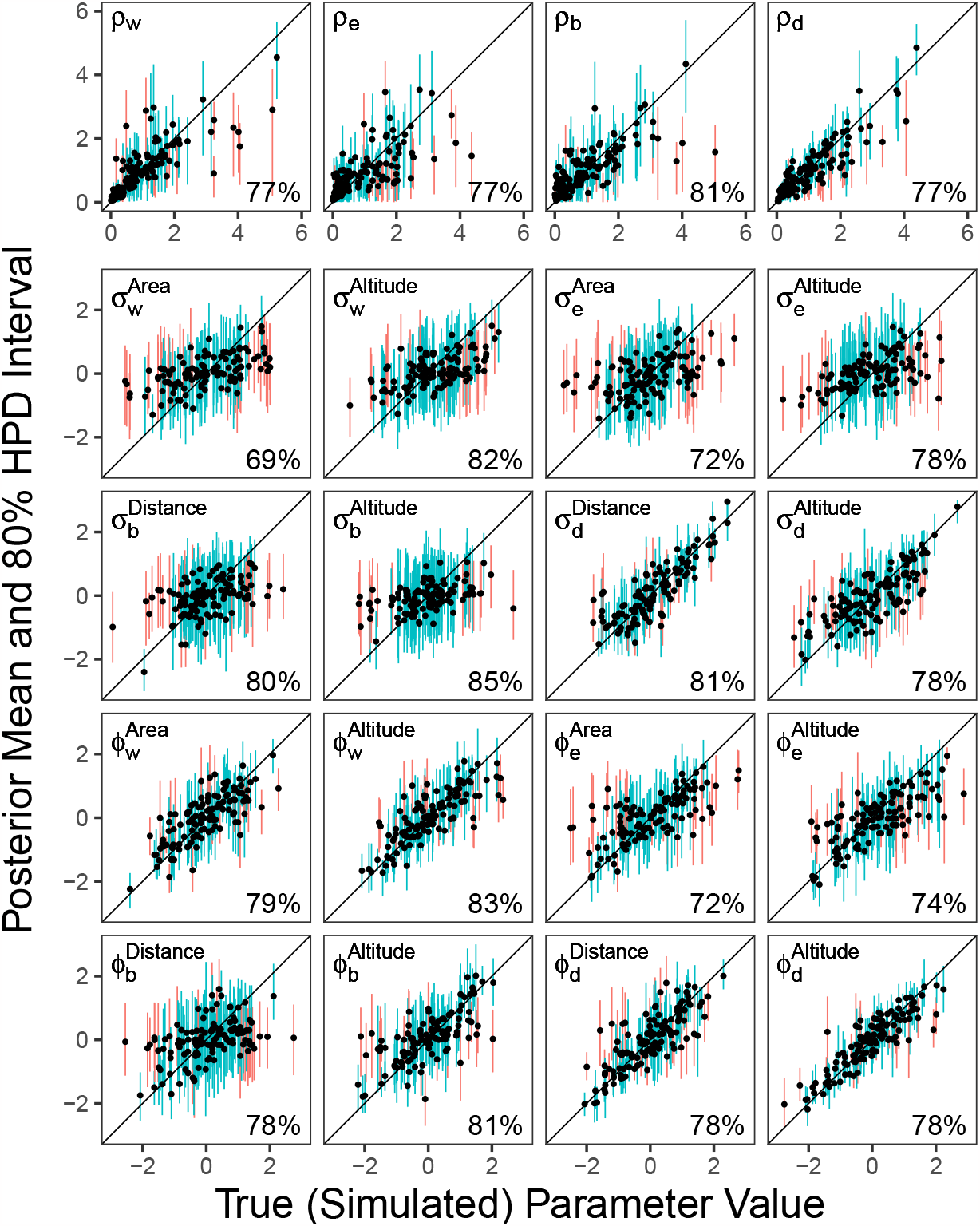
Quality of estimation and coverage of MultiFIG model parameters in simulations with five regions and medium tree size. Plots show true parameters on the *x* axis and estimated values on the *y* axis. Parameters are given in the upper left, and coverage percentages in the lower right of each plot. 80% HPD intervals which cover the truth are shown in blue, and 80% HPD intervals which do not cover the truth are shown in red. Some HPD intervals were clipped to the bounds of the plot (0 to 6 for *ρ* parameters, -3 to 3 for *σ* and *ϕ* parameters). Coverage analysis indicates that the inference method behaves appropriately, and plots indicate that most parameters can be inferred reasonably well (demonstrated by alignment with the 1:1 line).

In the analysis of the RJMCMC results (Figure S9), it is evident that RJMCMC does a better job assigning high probabilities when the effect parameter has a large (absolute) value, and is more variable in performance when the effect parameter has a low (absolute) value, presumably being confounded by other factors in absence of a strong feature effect.

### Liolaemus analysis

Here, we present posterior intervals from a traditional MCMC alongside RJMCMC posteriors for each parameter/process pairing (Figures S11, S12, S13 & S14). We also present joint RJMCMC probabilities for quantitative and categorical representations of the same features (Figure 6).

**Figure 6:**
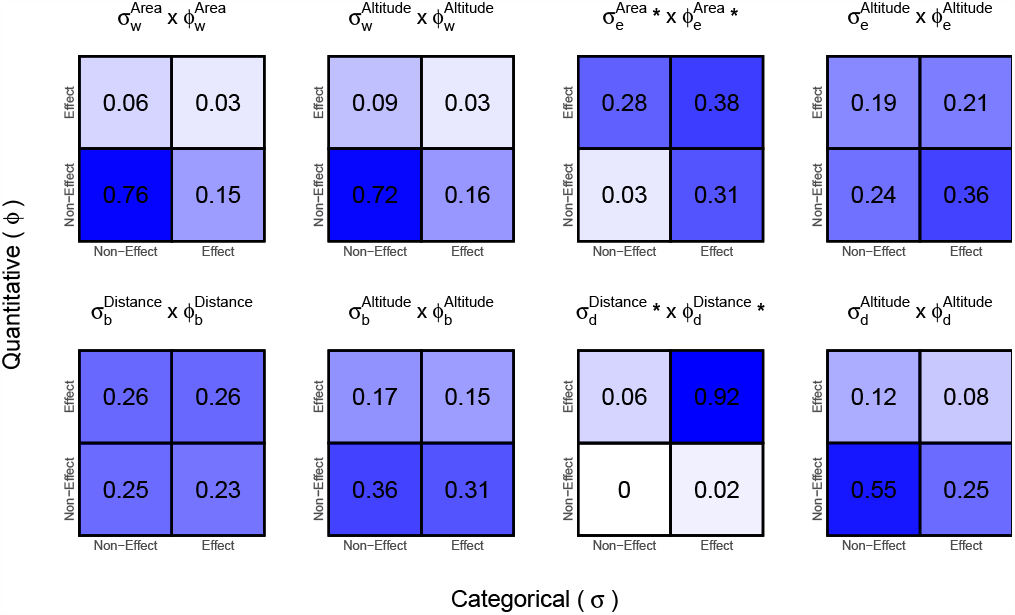
Joint probabilities of highly-correlated MultiFIG model parameters in the reversible-jump analysis of *Liolaemus*. The lower-left quadrant indicates the probability that a feature does not affect a certain process, since neither parameter associated with the feature (quantitative or categorical) shows an effect. The only feature/process pairings for which we can reject the null model (non-effect) are altitude/dispersal and area/extinction.

Of the 16 feature effect parameters examined, 4 were inferred to have a probable effect, uncovering several interesting relationships between features and processes. First, posterior densities indicate a probable negative relationship between geographical distance and dispersal. The results imply that both quantitative distance (mean distance between points in each region) and categorical distance (region adjacency) are important components of the model. RJMCMC posteriors also support this conclusion. Second, an analysis of joint RJMCMC probabilities of highly-correlated effect parameters (categorical and quantitative representations of the same feature) indicate that region area (size) is negatively correlated with rates of extinction. The model assigns a low probability (0.03) to the scenario where neither the quantitative or categorical parameter has an effect, meaning that region area likely relates to extinction in some capacity. However, the model does not assign a high probability of both parameters being simultaneously included (0.38), and does not seem to have a strong preference for either the quantitative or categorical representation.

The remaining 12 feature effect parameters were not inferred to have an effect. In these instances, MCMC and RJMCMC results were unable to reject the null (non-effect), and joint RJMCMC probabilities did not indicate the importance of either the quantitative or categorical representation of a particular feature. The model was also unable to assign *p >* 0.95 to any parameter being turned ‘off’, so we cannot conclusively determine that these features are not associated with biogeographic rates. Interestingly, one of the feature/process pairings where we did not find an effect was was altitude on within-region speciation. The parameters 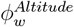 and 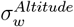 represent the impact of quantitative and categorical altitude on within-region speciation rates. The null effect (0) is well within the 95% posterior interval for both of the parameters, and the reversible jump MCMC supports a 0 value for both. Additionally, the joint RJMCMC analysis assigns highest probability to the scenario where neither parameter is incorporated (*p* = 0.72). Therefore, our model did not find any significant effect of quantitative or categorical altitude on within-region speciation.

Examining the posterior distribution for the number of ‘on’ parameters gives a more nuanced picture (Figure 7). Here, we can see over-representation of the 4 parameters 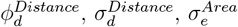, and 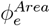 preferred by the individual/pairwise analyses. We can also see under-representation of some parameters (such as 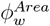 and 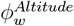), which indicates that the model prefers not to include these parameters. The model prefers scenarios with more than 4 parameters with probability *>* 0.95, but does not strongly suggest which parameters may be involved.

**Figure 7:**
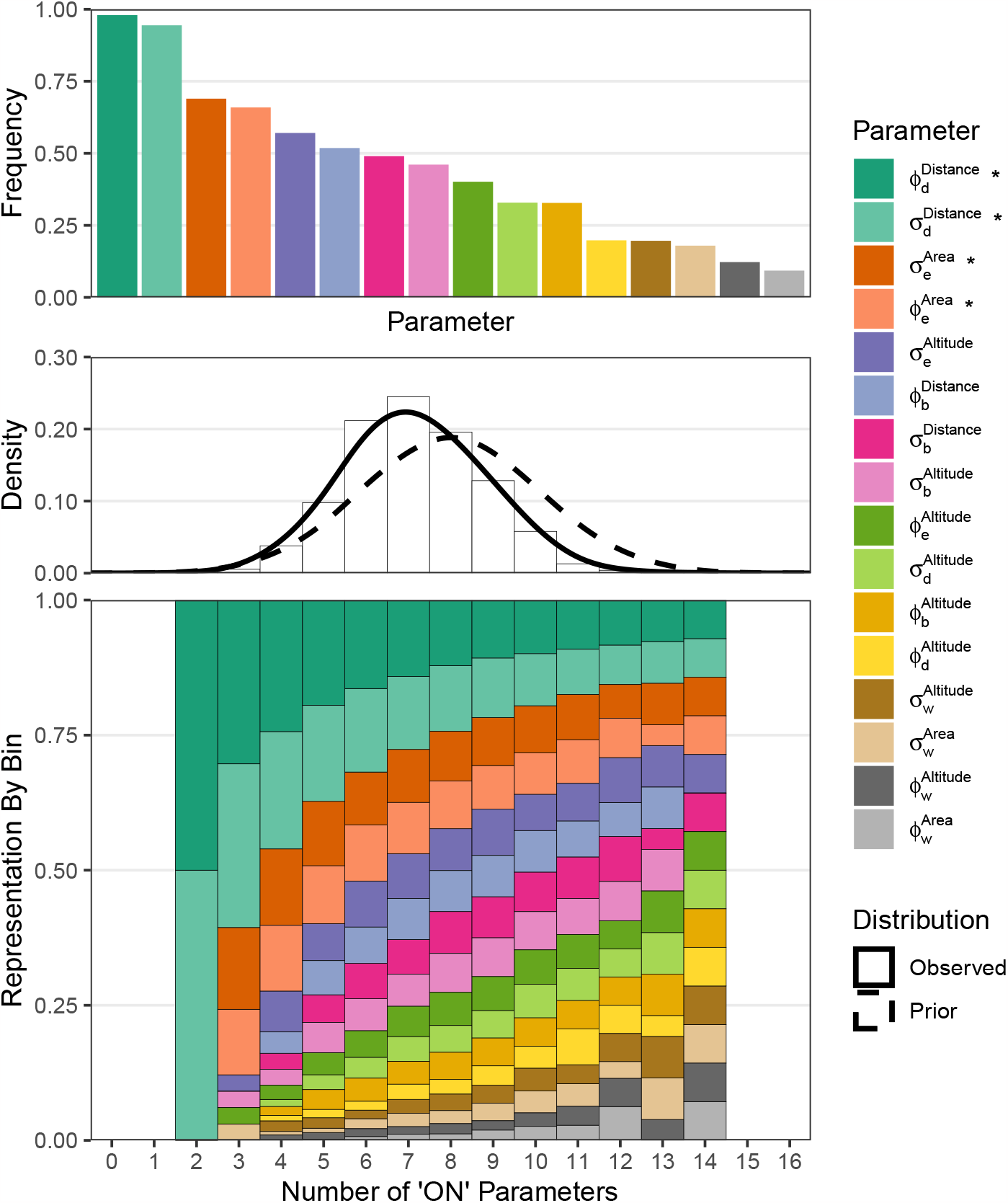
Summary of MultiFIG RJMCMC analysis over all considered feature-effect hypotheses. The top panel shows the frequency of being turned ‘on’ for each *ϕ* and *σ* parameter in the RJMCMC analysis. The other two panels show a more detailed view; the center panel shows the prior versus posterior distributions for the number of *ϕ* and *σ* parameters turned ‘on’ by RJMCMC, and the bottom panel shows the proportional representation of each parameter by bin. Parameters with inferred significance (from individual posteriors and joint reversible-jump probabilities) are marked with an asterisk.

## Discussion

Our study explored how MultiFIG can be used test and measure the many ways that regional features might shape corresponding biogeographic rates of dispersal, speciation, and extinction. Through simulation, we demonstrated the ability for MultiFIG to infer the effects of regional features on evolutionary processes. For the number of features examined (8 quantitative and 8 categorical), the model performed best in scenarios with more regions and larger trees (Landis et al. 2022). This is a sensible outcome, as we would expect an increase in the amount of data (more known feature values and more observed taxa) to improve parameter estimates. In all scenarios, the coverage analysis indicates that MCMC results can be trusted; if the 95% HPD interval ‘rules out’ a null effect, we can be reasonably confident that a feature is related to an evolutionary process. From the examination of RJMCMC analyses, we can see that RJMCMC is not a reliable tool for detecting effects under this model when used exclusively, particularly if those effects are small in magnitude. However, it is important to note that all parameters were turned ‘on’ during simulation (were given a non-zero value). If RJMCMC had the ability to perfectly discern which parameters did or did not have an effect, we might expect it to assign 100% probability to each parameter being ‘on’, but there are several reasons why it may not do this in practice. First, some parameters were assigned values very near zero during simulation, so the posterior surface may be too flat for RJMCMC to strongly prefer turning the parameter ‘on’. Second, even if an effect parameter has a large value, it may not actually impact evolutionary rates in a detectable way if the variance of a feature amongst regions or region pairs is small. If all regions had the same value for a particular feature, for example, it would be impossible to tell whether that feature was important, as there would be no point of comparison. Third, the simulated trees and biogeographies are drawn in a stochastic manner based on the rates provided, and contain a finite amount of data. Some trees may not perfectly reflect the underlying priors – for instance, small regions may have higher rates of extinction, but not every tree simulated under these conditions will show more extinction events in smaller regions. For these reasons (and others), we expect that RJMCMC will not assign 100% probability to every parameter being ‘on’. Importantly, the RJMCMC simulation results show that the production of strong false negatives is relatively rare, especially for parameters of large effect; if RJMCMC produces a strong probability of non-effect, we have good reason to believe the feature has little or no relationship with the process. This can be useful in conjunction with traditional posterior estimates and HPD intervals. Our RJMCMC simulation analysis did not investigate true negatives or false positives, since all parameters were turned ‘on’ in simulation. Further simulation studies should be conducted to determine the accuracy of assigning parameters to ‘on/off’ states using RJMCMC, incorporating simulations generated under smaller models with some parameters turned ‘off’.

Overall, the simulation study shows that MultiFIG has the ability to infer relationships between features and processes under a correctly-specified model. In this instance, it appears to perform well. Therefore, if we believe we have incorporated the features which are most closely related to evolutionary processes, and accept the assumption that all features act independently on evolutionary rates, we can be reasonably confident in our results. What happens when the model is misspecified remains to be studied. For instance, we may miss an important feature, or there may be interactions between features that we fail to capture, which can be modeled by treating those features as structured random variables (e.g. Landis 2017; Landis et al. 2021). It is important to reiterate that this model identifies *relationships*, but does not determine causality and certainly not direct causality; there may be other underlying causes for the relationships between features and processes, which could be teased apart using a hidden states framework (Beaulieu and O’Meara 2016; Caetano et al. 2018).

One interesting situation we considered was when two included features were highly correlated. This can sometimes lead to model misbehavior, masking the effects of one or both features. In our empirical analysis of *Liolaemus*, we examined the joint RJMCMC probabilities of non-effect for each pair of highly-correlated features (categorical and quantitative representations of the same information). This analysis is important; it allows us to incorporate multiple highly-correlated sets of parameters without losing inference capabilities. Through marginal probabilities alone, we were able to identify two feature parameters with small probabilities of non-effect: 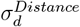 and 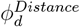. We were also able to examine the other features to determine whether marginal probabilities were being confounded by the duplication of information, and identified one additional feature with a low joint probability of non-effect: area with respect to extinction. It is worth comparing the two instances where an effect was inferred. In the first instance (distance), RJMCMC strongly preferred the incorporation of both parameters; in the other instance (size), RJMCMC preferred the incorporation of one parameter, but not necessarily both. While we cannot make any strong statement about the method’s consistency when it comes to selecting between models that represent a feature in a categorical or quantitative way (or both/neither), it is clear that the method does not always prefer more complex models (with multiple parameters representing similar regional feature) or simpler models (with only one parameter representing similar regional features).

Our model highlights two important feature/process relationships for *Liolaemus*: the relationship between distance and dispersal, and the relationship between area and extinction. A positive value for 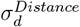 and a negative value for 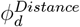 indicate that adjacent, closer-together regions have higher rates of dispersal. A negative value for 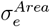 and a negative value for 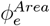 indicate that smaller regions have higher rates of extinction. These relationships are sensible, as distance (Landis et al. 2022) and area-size (Rabosky and Glor 2010; Thomson et al. 2021; Quintero et al. 2023) have previously been implicated in diversification dynamics in other reptile systems using SSE models.

Interestingly, one of the feature/process pairings where our model did not infer significance for either *σ* or *ϕ* was altitude and within-region speciation. It was our initial hypothesis that one (or both) of the altitude parameters would show an effect on speciation, since Andean regions possessed higher rates of speciation in the analyses of Esquerré et al. (2019), which examined the family Liolaemidae (including the *Liolaemus* genus). Abiotic (e.g. climatic) and biotic (e.g. competitive) factors are expected to differently shape the ecological niches of otherwise similar lowland versus highland species (Lomolino 2001), which should create different ecological opportunities for speciation and risks for extinction. The outsized historical effects cyclical climate change would have on mountains should further exaggerate differences in net diversification (Flantua et al. 2019; Mastretta-Yanes et al. 2015). That said, our categorization of highland/lowland regions did not initially align with the Andean/non-Andean classification of Esquerré et al.. In our analysis, Patagonia is classified as a ‘lowland’ region, since its mean altitude is below the mean for all regions, but Esquerré et al. classify Patagonia as an Andean region, as the southern part of the Andes falls within Patagonia. Patagonia was thought to be an important source of speciation in the Esquerré et al. analysis. Therefore, we performed a second analysis, replacing our highland/lowland classification with an Andean/non-Andean classification for all six regions following Esquerré et al. (2019). Even with this classification system, our model did not infer significance for parameters associating Andean/non-Andean status with rates of speciation. However, when we compared relative rates of within-region speciation among regions, Andean regions (in both highland/lowland and Andean/non-Andean analyses) had higher within-region speciation rates, even if the model did not indicate that the parameters associated with altitude were significantly different from zero. We also performed an analysis with a reduced set of features that only assigned regions an Andean or non-Andean classification, but did not incorporate areas, distances, or quantitative altitudes. This analysis is more closely related to the two-state GeoSSE model that Esquerré et al. used to infer higher rates of speciation in Andean regions, but still utilizes six regions. It similarly supported higher rates of speciation in Andean regions (1.28 times higher in Andean versus non-Andean regions), although parameters associated with this relationship did not show significance, much like in the full analyses with all sixteen parameters.

The posterior for the preferred number of parameters offers additional insight into the complex relationships between region features and biogeographic rates. Models with only 4 parameters were not ultimately preferred, even when no additional parameters could be individually selected. One explanation is that there may be rate variation in the system that is unrelated to the parameters we chose. Additionally, rate variation could be created by some complex interactions of our features which we did not examine, or the model may lack the ability to discern between some correlated parameters (we only examined certain 2-parameter joint probabilities). It is also possible that a different choice of priors would lead the model to identify additional significant parameters.

These results highlight the importance of incorporating multiple regional features, both categorical and quantitative, into inferences of biogeographic rates. In addition to recovering differences in speciation rates between Andean and non-Andean regions, we were able to identify important distance/dispersal and area/extinction relationships, which could not have been investigated in a two-region GeoSSE framework. The MultiFIG model can be extended to include several features that are expected to have an affect on different processes, and help determine which of those features actually possess a strong link to particular processes. The model can also be expanded to include additional features, or hypothesized relationships amongst feature effects.

Although many regional features change over time, we should expect historical rates of biogeographic processes might also change in kind. However, the FIG model currently makes use of only present-day regional feature data. When analyzing clades of extant taxa, previous work from our collaborations has found that ignoring paleogeography often has little effect on simpler biogeographic parameter estimation or model selection tasks (Landis 2017; Landis et al. 2021). We suspect this is because the highest proportion statistical information for powering models comes from living species and the regions they currently inhabit, with little coming from historical inferences phylogenetically extrapolated from those extant taxa. But we believe the reassurances end there because, not surprisingly, we have repeatedly found that ignoring or mischaracterizing paleogeography misleads divergence time and ancestral state estimates buried deeper in the tree (Landis 2017; Landis et al. 2018, 2021). To address this gap, future work will expand the FIG framework to allow for regional features to vary over time, enabling it to model a wider range of hypotheses concerning historical relationships between regional features and biogeographic rates, evolving biome affinities among lineages (Crisp et al. 2009; Lichter-Marck and Baldwin 2023), fossil taxon occurrence and preservation (Xing et al. 2016; Saupe et al. 2019; Landis 2021), and phylogenetic divergence times (Landis 2021).

## Funding

This study was supported by the National Science Foundation (NSF) through the Graduate Research Fellowship Program awarded to Sarah Swiston (NSF DGE 2139839) and through a research grant awarded to Michael Landis (NSF DEB 2040347). Any opinions, findings, and conclusions or recommendations expressed in this material are those of the author(s) and do not necessarily reflect the views of the National Science Foundation.

## Acknowledgements

We are grateful to members of the Landis and Myers labs at Washington University in Saint Louis and the Tello lab at the Missouri Botanical Garden for feedback. This work was also enriched by ongoing collaborations with Drs. Isaac Lichter-Marck and Felipe Zapata at UCLA.

---

**Figure S8:**
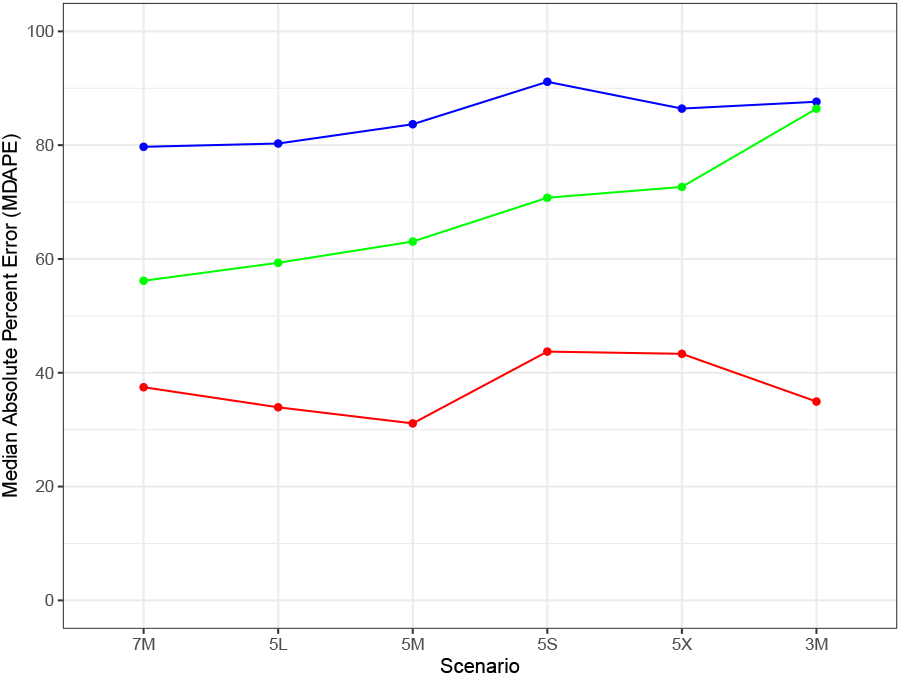
Median absolute percent errors for each set of parameters (*ρ, ϕ*, and *σ*) grouped by number of regions and tree size. Extra small (X): 25–50 taxa, small (S): 51–100 taxa, medium (M): 101–200 taxa, large (L): 201–350 taxa. Scenarios are loosely ordered from information-rich (left) to information-poor (right). In general, estimates were poorer in scenarios with smaller trees and fewer regions.

## Supplementary Materials

**Figure S9:**
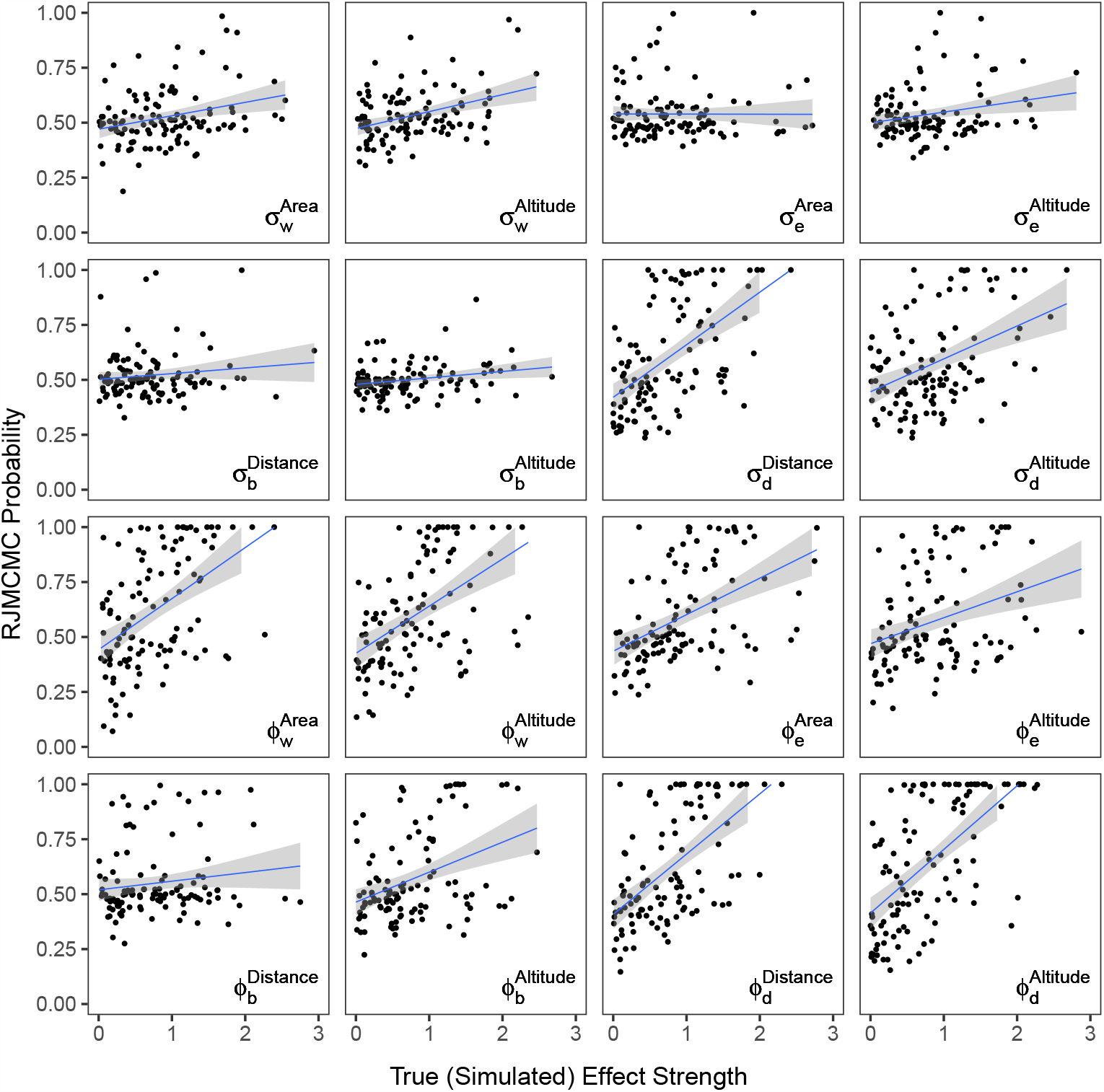
RJMCMC probabilities plotted against parameter strength (the absolute value of the true value of the effect parameter) in simulations with five regions and medium tree size. Most parameters show a moderate probability of being ‘on’, which is expected given that all simulations included a non-zero effect strength for each parameter. Overall, ‘Stronger’ parameters (with effect values farther from zero) show a higher RJMCMC probability of being ‘on’.

**Figure S10:**
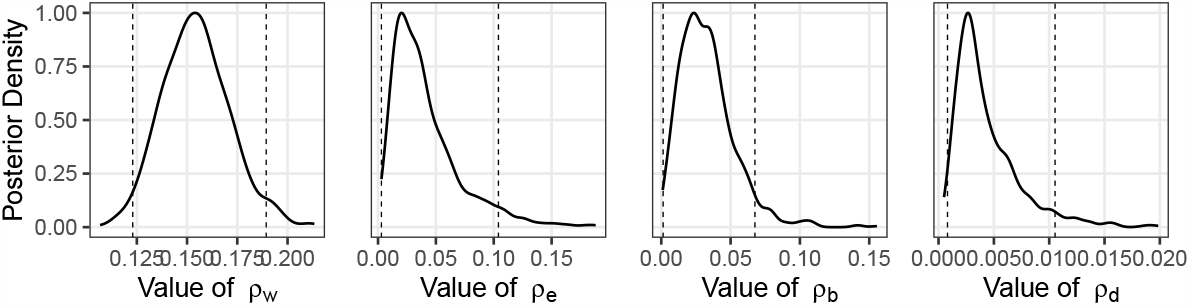
Posterior distributions for estimated base rates of within-region speciation [*ρ*_*w*_], extinction [*ρ*_*e*_], between-region speciation [*ρ*_*b*_], and dispersal [*ρ*_*d*_]. Dashed lines represent 95% HPD intervals.

**Figure S11:**
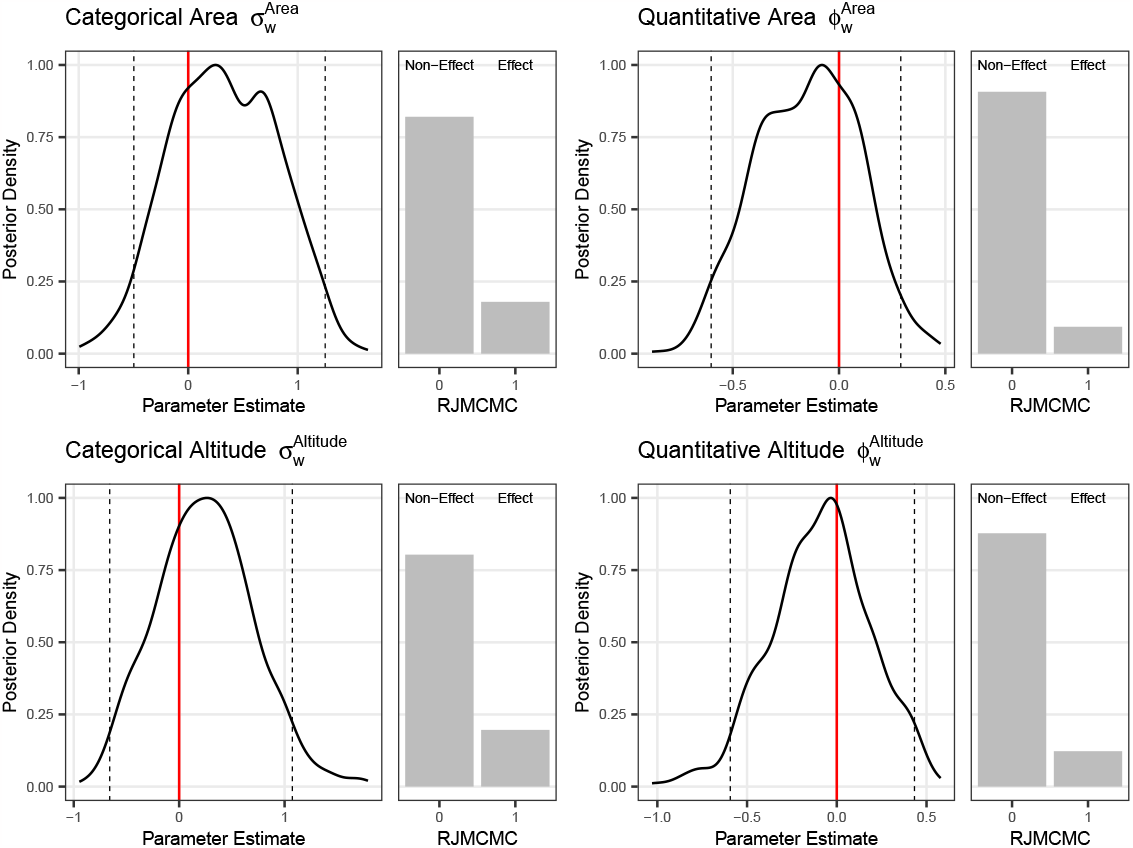
Impact of features on within-region speciation. Included are posterior distributions (left) and RJMCMC probabilities (right). Dashed lines represent 95% HPD intervals.

**Figure S12:**
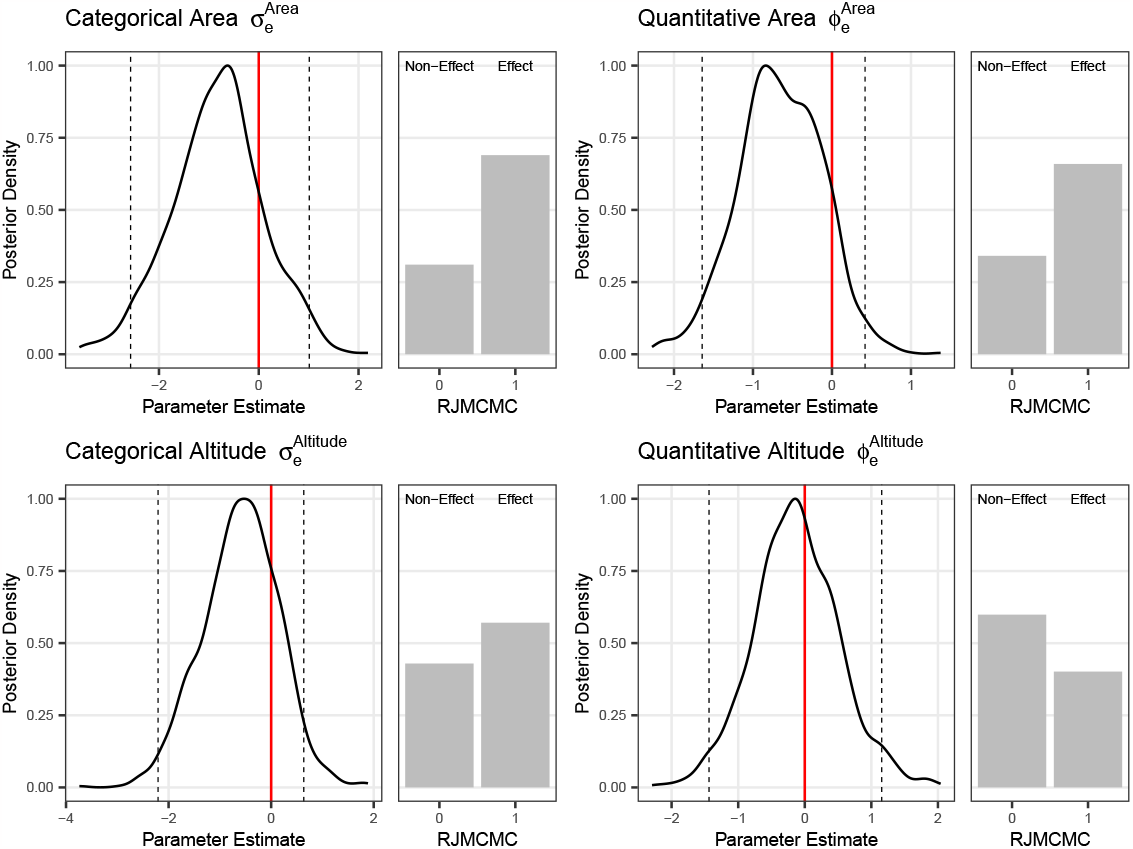
Impact of features on extinction. Included are posterior distributions (left) and RJMCMC probabilities (right). Dashed lines represent 95% HPD intervals.

**Figure S13:**
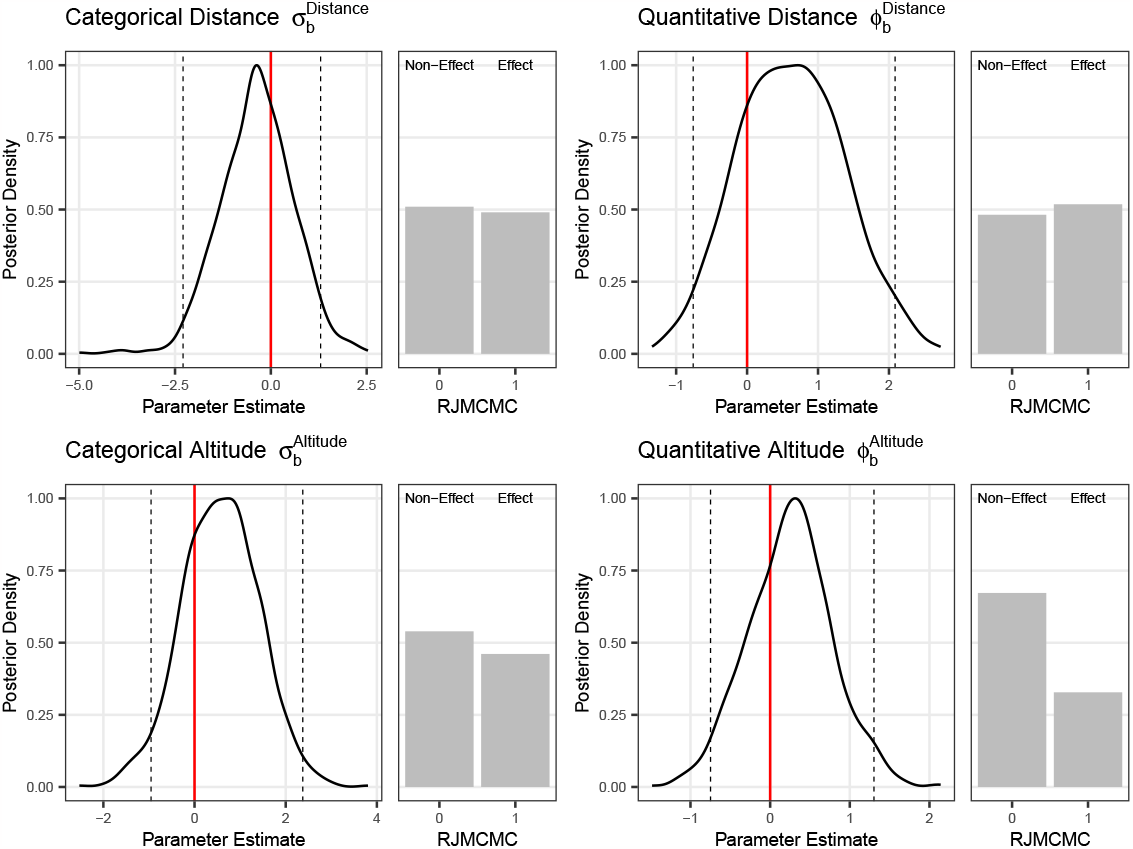
Impact of features on between-region speciation. Included are posterior distributions (left) and RJMCMC probabilities (right). Dashed lines represent 95% HPD intervals.

**Figure S14:**
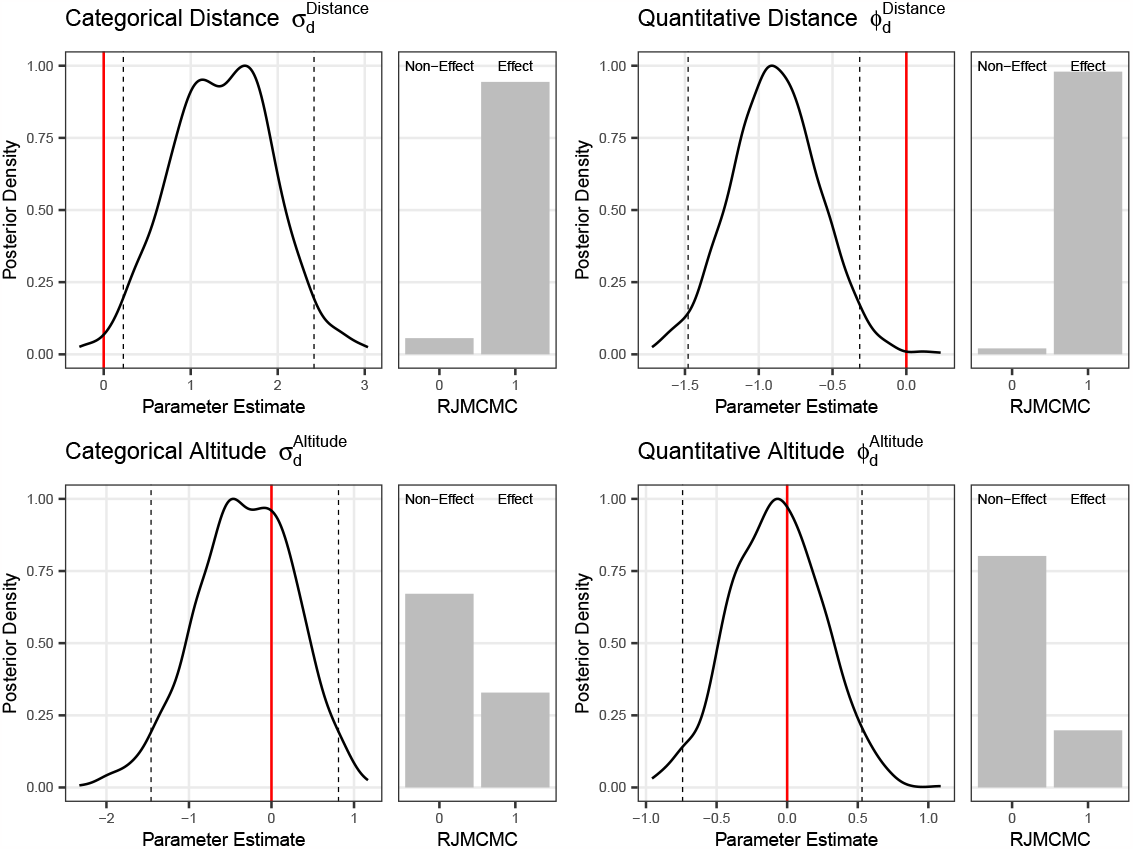
Impact of features on dispersal. Included are posterior distributions (left) and RJMCMC probabilities (right). Dashed lines represent 95% HPD intervals.

**Figure S15:**
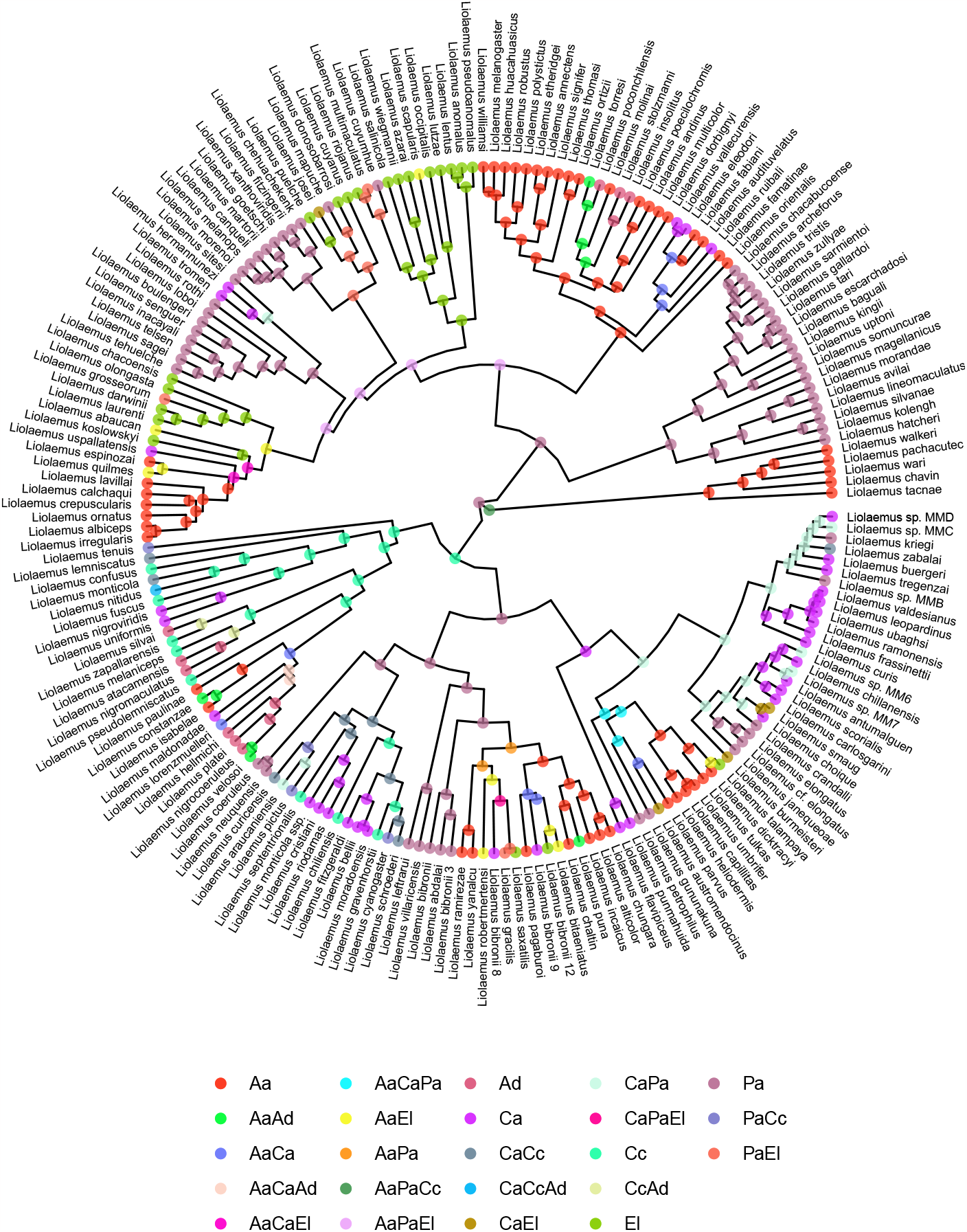
Ancestral state reconstruction for *Liolaemus*. Highest-probability states are recorded at internal nodes. Important abbreviations: Aa (Altiplanic Andes), Ca (Central Andes), Pa (Patagonia), Cc (Central Chile), Ad (Atacama Desert), El (Eastern Lowlands). States may include multiple regions.

**Table S2:** 80% HPD coverages of each model parameter grouped by number of regions and tree size. Extra small (X): 25–50 taxa, small (S): 51–100 taxa, medium (M): 101–200 taxa, large (L): 201–350 taxa. No model parameters or simulation scenarios demonstrate especially poor coverage, and the inference method appears to behave appropriately.

| Parameter | 3M | 5X | 5S | 5M | 5L | 7M |
| --- | --- | --- | --- | --- | --- | --- |
| $\rho_w$ | 80 | 80 | 81 | 77 | 70 | 73 |
| $\rho_e$ | 77 | 82 | 76 | 77 | 79 | 70 |
| $\rho_b$ | 81 | 79 | 84 | 81 | 77 | 83 |
| $\rho_d$ | 83 | 80 | 74 | 77 | 82 | 70 |
| $\sigma_w^{Area}$ | 80 | 86 | 78 | 69 | 74 | 88 |
| $\sigma_w^{Altitude}$ | 72 | 78 | 69 | 82 | 75 | 78 |
| $\sigma_e^{Area}$ | 80 | 77 | 80 | 73 | 80 | 75 |
| $\sigma_e^{Altitude}$ | 75 | 77 | 81 | 78 | 77 | 71 |
| $\sigma_b^{Distance}$ | 78 | 74 | 81 | 80 | 77 | 75 |
| $\sigma_b^{Altitude}$ | 76 | 83 | 80 | 85 | 74 | 81 |
| $\sigma_d^{Distance}$ | 77 | 78 | 71 | 81 | 80 | 72 |
| $\sigma_d^{Altitude}$ | 78 | 77 | 78 | 78 | 77 | 78 |
| $\phi_w^{Area}$ | 72 | 77 | 77 | 79 | 73 | 76 |
| $\phi_w^{Altitude}$ | 79 | 75 | 77 | 83 | 75 | 69 |
| $\phi_e^{Area}$ | 72 | 73 | 75 | 73 | 73 | 76 |
| $\phi_e^{Altitude}$ | 71 | 74 | 84 | 74 | 76 | 76 |
| $\phi_b^{Distance}$ | 73 | 77 | 78 | 78 | 78 | 75 |
| $\phi_b^{Altitude}$ | 71 | 78 | 81 | 81 | 74 | 84 |
| $\phi_d^{Distance}$ | 83 | 70 | 82 | 78 | 82 | 76 |
| $\phi_d^{Altitude}$ | 78 | 74 | 86 | 78 | 77 | 75 |

## Notes

https://bitbucket.org/sswiston/multifig/src/master/

https://hub.docker.com/r/sswiston/rb_tp

https://revbayes.github.io/tutorials/multifig.html

### Competing Interest Statement

The authors have declared no competing interest.

